# Estimating Residue-Specific Accuracies of Protein Structure Models Using Molecular Dynamics Simulations

**DOI:** 10.1101/439760

**Authors:** Jing-Hao Hu, Sang-Ni Xun, Hao-Nan Wu, Yun-Dong Wu, Fan Jiang

**Affiliations:** Laboratory of Computational Chemistry and Drug Design, State Key Laboratory of Chemical Oncogenomics, Peking University Shenzhen Graduate School, Shenzhen, 518055, China.; College of Chemistry and Molecular Engineering, Peking University, Beijing, 100871, China.

**Author notes:** Correspondence and requests for materials should be addressed. (F.J.).

## Abstract

Estimating the accuracy of a structure model is very crucial to promote the usefulness of protein structure prediction methods. Currently, a vast majority of successful model quality assessment (or model accuracy estimation, MAE) methods are knowledge-based. Based on molecular dynamics (MD) simulation with a recently developed residue-specific force field (RSFF2), we develop a method for absolute MAE at per-residue level. Using a training set of 31 models and a test set of 24 models from different proteins, the MAE performance of our MD-based method can reach or even exceed state-of-the-art single-model MAE methods within a short simulation time (less than one nanosecond). In addition, a simple combination of knowledge-based method with the MD-based method can give more accurate MAE than any of the constituent methods.

## Introduction

Recently, the cost of obtaining new protein sequences has decreased dramatically due to significant progresses in high-throughput DNA sequencing technology.^1, 2^ On the other hand, to obtain high-resolution three-dimensional (3D) structures experimentally is still quite expensive, despite numerous effects. Therefore, there is a huge and increasing gap between the numbers of known structures and sequences.^3-5^ Due to the importance of protein structures in modern biological research and drug discovery,^6-8^ protein structure prediction (PSP) has become a cost-effective and high-throughput way to provide structure information for biological and pharmaceutical studies.^9, 10^

There are various methods for PSP^11-13^, including homology modelling,^14^ threading,^15^ fragment assembly,^16, 17^ coevolution-based analysis,^12, 18^ and *ab initio* folding.^19, 20^ The predicted protein models can span a broad range of accuracies and are suitable for different applications.^7^ They usually have lower qualities compared with experimental structures, significantly limiting their usefulness especially for certain important applications including structure-based drug design.^21-24^ Besides, a single PSP method can often generate a number of structure models for a given protein target, whereas it is very difficult to pick out the most accurate one(s).^25^ The discrimination of accurate models from decoys has been a key factor to further increase the accuracy of PSP.^26^ Thus, a reliable method of ranking models is urgently needed, which we called *relative* model accuracy estimation.^26, 27^

Before using a structure model, it can be very beneficial to check whether it is accurate enough to avoid the risk of misuse. Thus, there is also a need for *absolute* model accuracy estimation (or simply called MAE) to estimate the discrepancy between a model structure and the (unknown) actual native structure, which can be performed on global or local level. The *global* MAE evaluates the overall quality of a model, while the *local* MAE estimates the error at residue-specific or atomic level. Basically, local MAE is more useful because that global score can often be inferred from summarizing local scores. Moreover, it can provide accuracy information of different regions, which can be used as a guide for further refinement.^26^

Some early MAE tools can evaluate a protein model by checking the correctness of stereochemistry, such as WHAT-CHECK,^28^ PROCHECK,^29^ and MolProbity.^30^ Besides, many knowledge-based potentials derived from statistical analysis of features in known protein structures can discriminate near-native structures from decoys (relative MAE).^31^,^32^ Recently, there are developments of absolute MAE methods such as the ProQ series^26, 33-35^ using supervised machine learning on multiple features including evolutionary information, residue environment compatibility or components from knowledge-based potentials. Besides these *single-model* methods, there are also consensus-based (clustering) methods, in which the quality prediction of a model is correlated to its similarity to other models, such as Pcons.^36^ Recently, hybrid methods that combine single-model and consensus-based approaches have also been developed.^37, 38^

The MAE was introduced into the Critical Assessment of protein Structure Prediction (CASP) experiment as an independent category since 2006, traditionally called model quality assessment (MQA).^39-44^ As far as we know, a vast majority of successful MAE methods are knowledge-based.^45^ However, with rapid increasing of computing power, the physics-based methods have shown promising future for PSP. Especially, molecule dynamics (MD) simulation, as a useful tool for studying protein dynamics, have been successfully applied in *ab initio* protein folding^19, 20^ and structure refinement.^46-49^ The successful use of MD simulation in MAE has not been reported, but we think that it may work based on a physically sound assumption: native structure (free energy minimum) of a protein can be well kept during a MD simulation, but a non-native structure is less stable and will deviate from the initial coordinates.

Here, we evaluate the applicability of MD simulations in single-model absolute MAE at residue level. Our recently developed residue-specific force field RSFF2^50^ was used, which is an improvement of AMBER ff99SB^51^ by fitting conformational distributions from PDB coil library using residue-specific torsional parameters.^52-54^ Previous studies have shown that RSFF2 can well reproduce conformational behaviors of peptides and proteins,^55^ and predict the crystal structures of cyclic peptides accurately.^20^ We first performed MD simulations of a series of target models to find suitable patterns that are related to residue-level model quality. Then, we compared our MD-based MAE method with other popular single-model methods and investigated whether a combination of MD-based and knowledge-based methods can achieve better performance. Finally, we further analyzed the MD trajectories to explain our findings and make connections to MD-based model refinement.

## Material and Methods

### Protein training and test datasets

Thirty-one single-domain models (targets) from the refinement category of CASP 8-10, excluding those with missing residues in corresponding experimental structures, were chosen as the training dataset (Table S1). Different models corresponds to different proteins, with their sizes ranging from 63 to 235 residues. They have diverse secondary structures and a large coverage range (1.3 −7.7 Å) of Cα-RMSD to corresponding experimental structures. Meanwhile, twenty-four targets from the refinement category of CASP 11 were chosen as the test dataset. All structures were downloaded from the CASP website.

### Molecular dynamics (MD) simulations

All MD simulations were carried out using the *Gromacs* 4.5.4 software^51, 55^ with the RSFF2 force field.^50^ The hydrogen atoms in each model were added by *Gromacs* under neutral pH. Each protein was solvated in an octahedron box with the TIP3P water model. The minimum distance between the protein and the box boundary was 4.5 Å. Na+ or Cl-ions were added to neutralize the protein charges and to achieve an ion concentration of 0.05 mol/L. The electrostatic interactions were calculated by the particle-mesh Ewald method with real-space cutoff of 9 Å. The cutoffs of van der Waals interactions were also 9 Å, with long-range dispersion correction. The integration time step was 3 fs, with all bonds involving hydrogen atoms constrained using LINCS. The Berendsen method was used to maintain temperature and pressure (1 atm). For each target, after a 5000-step energy minimization, a 200 ps NPT simulation was performed with temperature increasing from 10 to 298 K within the first 40 ps, to equilibrium the solvent and box volume. During this process, positions of all Cα atoms were restrained by harmonic potentials of 10 kJ/mol/Å^2^. Then, parallel NVT productive simulations were performed from different initial velocities. Structures were stored every 1.5 ps during the simulations.

## Results

### Structural deviations in MD correlate well with actual model errors at residue level

MD simulations of thirty-one protein models (training dataset) were performed. For each protein at each of the five temperatures (330K, 360K, 380K, 400K and 430K), five parallel MD simulations were carried out for 3 ns. As shown in Figure 1a and 1b for two representative cases, residue-level root-mean-square deviations (RMSD in time-average) from MD simulations (eq. 1) correlate significantly with corresponding actual residue-level modeling errors (eq. 2). When considering all residues in the 31 models together, the Pearson’s correlation coefficient *r* between MD-derived RMSD values and actual modeling errors (0.64, Figure 1c) is similar to that from a state-of-the-art MAE method ProQ3D^35^ based on deep learning (*r* = 0.62, Figure 1d).

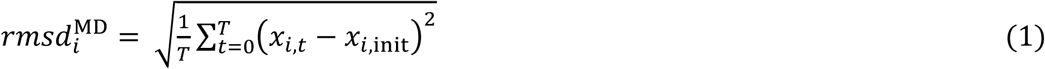

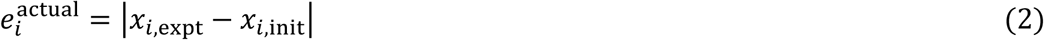

**Figure 1.**
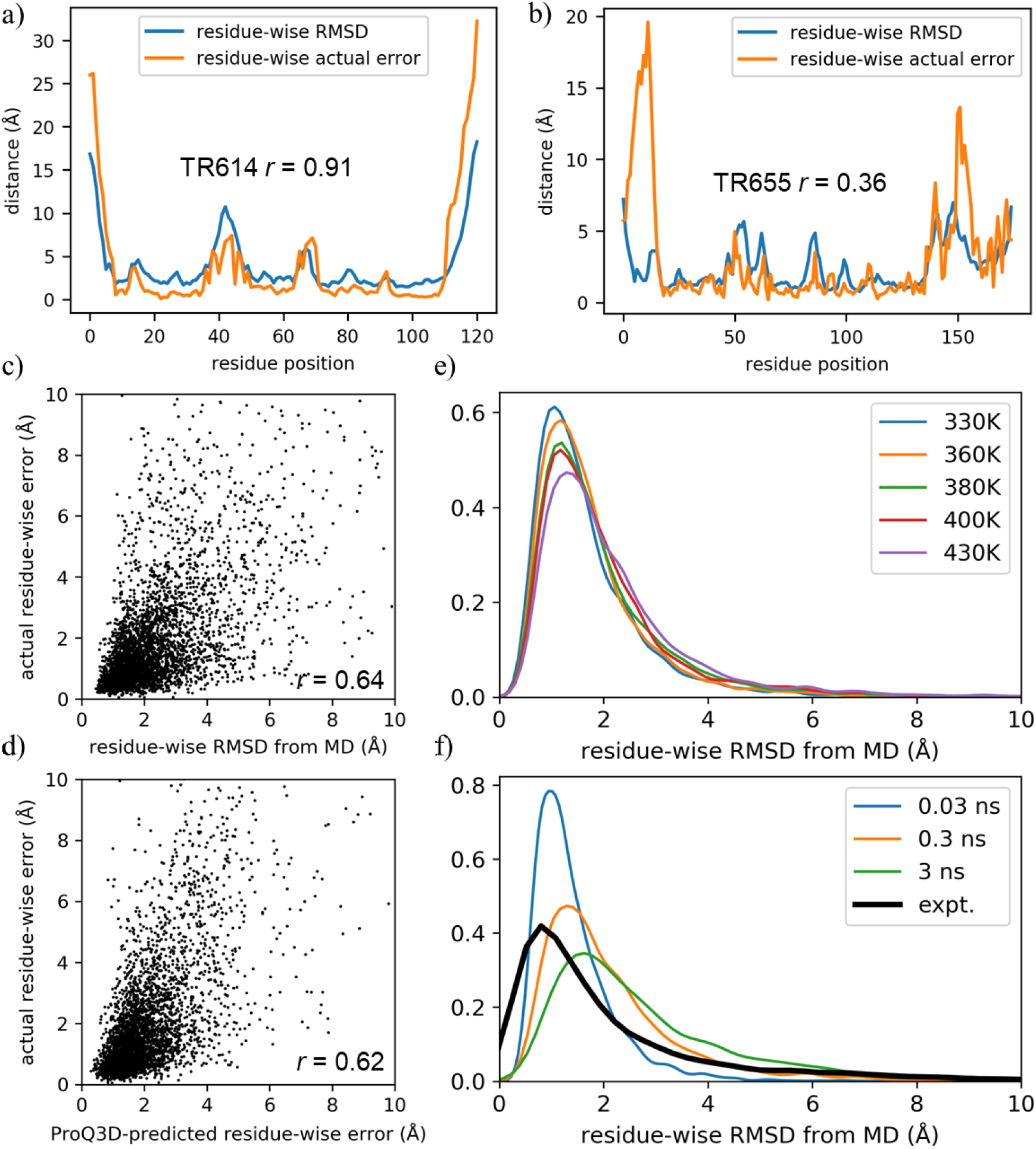
a) and b) Actual modeling error (defined in eq. 2) and time-averaged RMSD (defined eq. 1) from MD simulation (380K, 0.3 ns) at each residue position for protein target TR614 or target TR655, Pearson’s correlation coefficient *r* was also given for each case. c) and d) Actual modeling errors plotted against the MD-derived RMSD values (c) or the ProQ3D predictions (d) for all residues in the training set (31 proteins). e) and f) Residue-wise RMSD distribution at different simulation temperatures (c) and simulated times (d). The bold black line shows the distribution of actual residue-wise errors.

To apply eq. 1, all structure frames in a MD trajectory was optimally superposed to the initial model. Here, *x*_*i*,init_ and *x*_*i,t*_ are Cα positions of the *i*^th^ residue at simulation time *t*=0 (the initial model) and *t*=*t*, respectively. *T* is the trajectory length (number of frames) used for the calculation. Please note that this residue-wise RMSD is not to measure the fluctuation (flexibility) of each residue around its average position as the RMSF analysis commonly used in MD studies, but to measure the average *deviation* from the initial position of each residue. In eq. 2, *x*_*i*,expt_ is Cα position of the *i*^th^ residue in the experimental structure. The distance was also calculated after optimal superposition.

Interestingly, different simulation temperatures (330K – 430K) give quite similar residue-wise RMSD distributions (Figure 1e), with slightly larger average RMSD values and more dispersed distributions from higher temperatures. As expected, longer simulation times make RMSD distributions shift to larger values (Figure 1f). MD simulations of 0.03 ns can give a RMSD distribution with peak position similar to the actual error distribution, but it is narrower. We cannot find a simulation time and temperature that can give MD-derived RMSD distribution similar to the actual error distribution, indicating that certain kind of normalization is needed to make absolute MAE prediction.

### From MD-derived RMSD to absolute local MAE predictor

Here, we use a simple linear transformation to calculate the MD-derived error estimates:

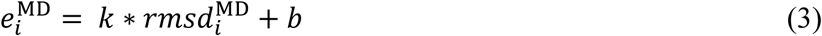

The coefficients *k* and *b* were calculated based on the following statistics of MD-derived RMSD (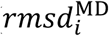, eq. 1) and actual errors (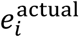, eq. 2) in our training dataset:

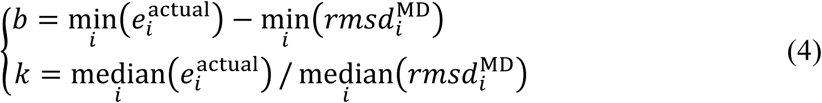

Obviously, *k* and *b* depend on the simulation time (Figure 2a). As shown in Figure 2b, the normalized 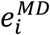 have similar distributions under different simulation times, and are in good agreement with the target distribution 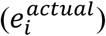.

**Figure 2.**
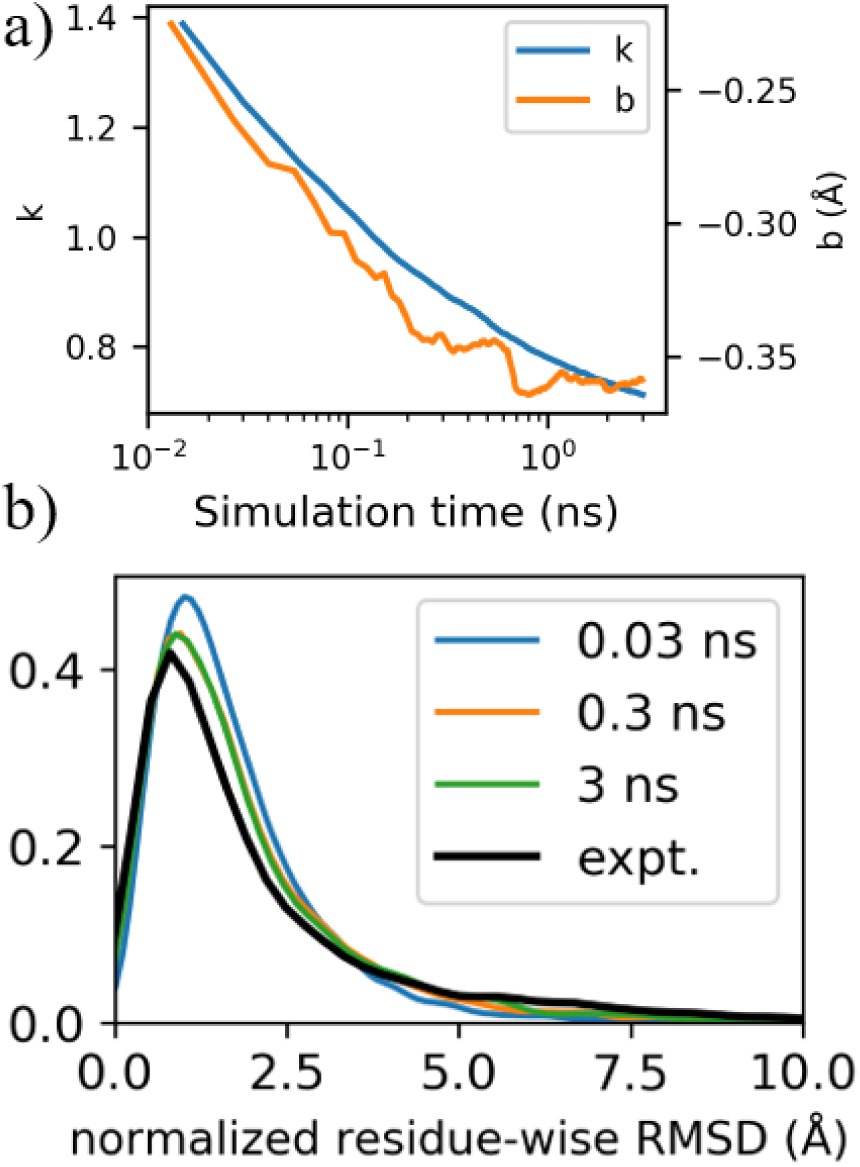
a) Normalization parameters *k* and *b* (eq. 3) over simulation time (in logarithmic coordinates), from MD simulations at 380K. b) Distributions of the normalized residue-wise RMSD (MD-derived error estimator) under different simulation times. The bold black line shows the distribution of actual residue-wise errors.

To better evaluate the performance of our MAE predictor, the distance measures 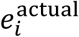 and 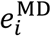 were converted to s-scores in the [0, 1] range:^56^

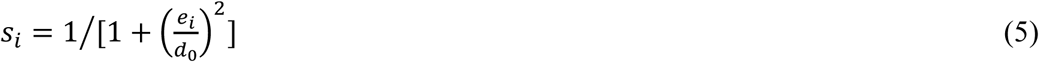

where *e*_*i*_ is the position error of the *i*^th^ residue, *d*_0_ is set to 3 Å in consistent with MAE methods of ProQ series. Higher s-score indicates higher model accuracy. Understandably, using s-score for MAE is better than direct use of distances. For poor quality parts in a protein model, the residue-specific RMSD can change in a fairly large range (e.g. 5 ∼ 15 Å) without too much difference in accuracy, whereas a smaller change of RMSD from 1.5 Å to 3 Å can significantly affect applicability of a model. Figure 3 shows the distributions before and after distances were converted into s-scores, and clearly most residues in our dataset have relatively high s-scores.

**Figure 3.**
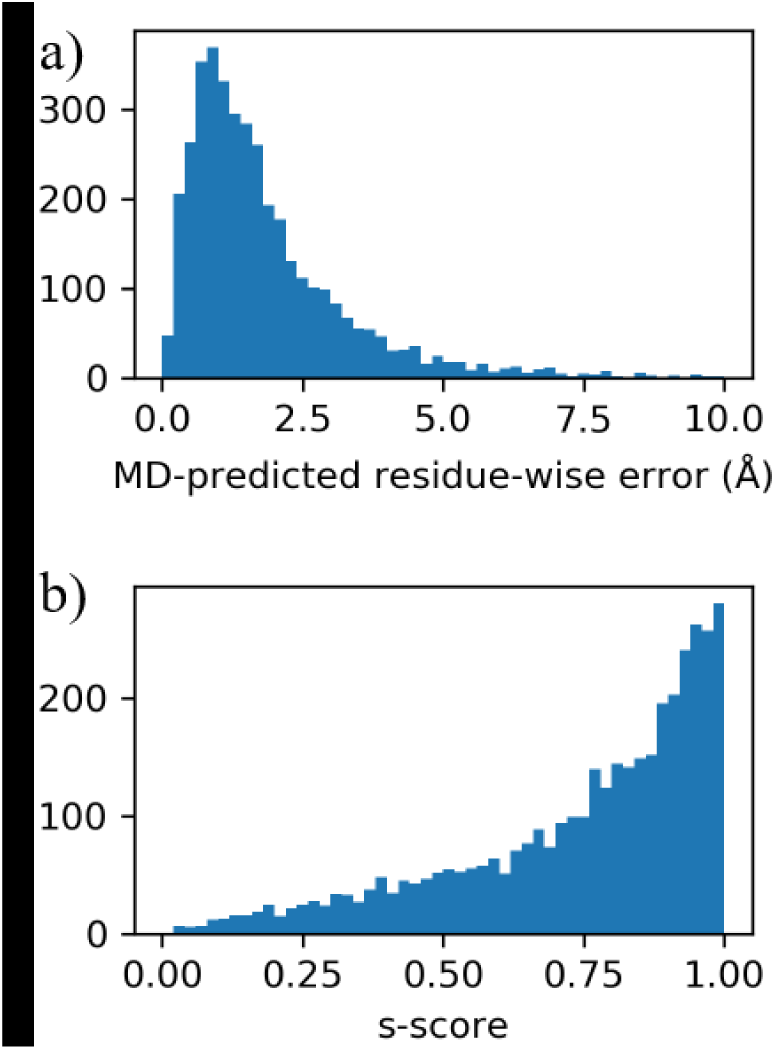
Distribution of MD-predicted residue-wise errors in histograms: a) before and b) after distances are converted into s-scores. Results from 380K MD simulations (0.3 ns) were shown.

### Comparison with state-of-the-art methods

During the first 0.3 ns of MD simulations, the accuracy of residue-specific MAE predictions (as measured by the correlation *r* between MD-predicted s-scores 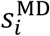 and real s-scores 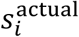) increases quickly to be higher than ProQ3^33^ predictions (Figure 4). After 0.5 ns of MD simulations, the results from the five parallel MD trajectories did not change significantly, which are better than ProQ3 and slighter worse than ProQ3D. However, when the residue-wise RMSD values from five individual trajectories were averaged before converted to s-scores, performance better than ProQ3D can be achieved. Correlation coefficients *r* from MD-based predictor reach 0.66 within 0.3 ns and finally reach 0.69 within 3 ns, whereas ProQ3D gives *r* of 0.64.

**Figure 4.**
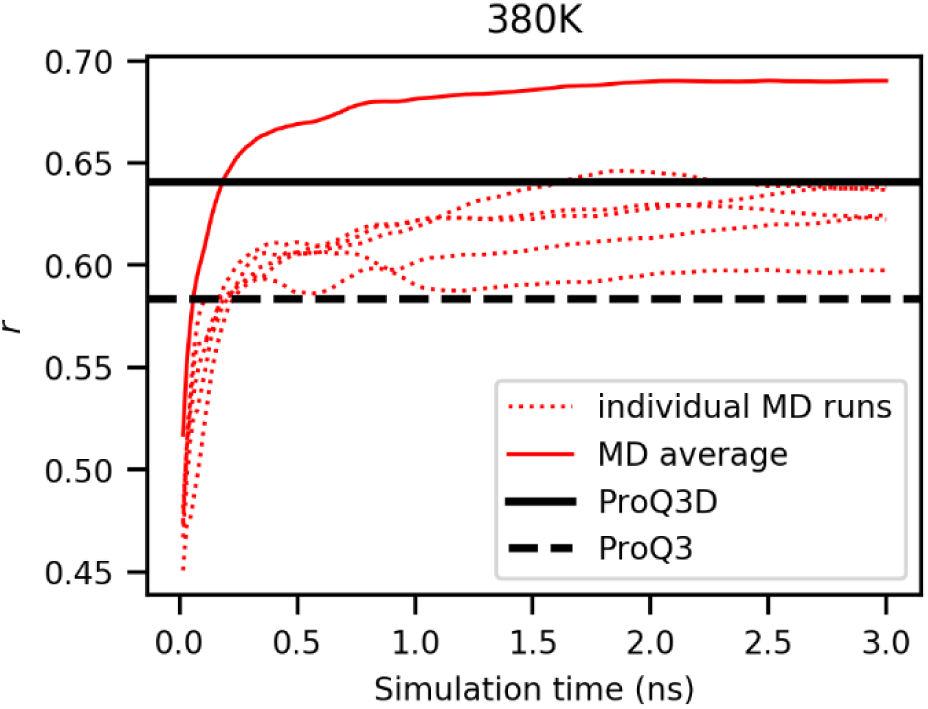
The correlation coefficient (*r*) between MD predictions and actual errors (both in s-scores) in the training dataset over 3 ns simulation time at 380K. The dash red lines are the predictions from five individual MD runs, and the solid red line is the average of the five MD runs’ predictions. The results of ProQ3 and ProQ3D are shown in the horizontal black dash line and black solid line, respectively.

In the local MAE (MQA) competition of the CASP, the Accuracy of Self-Estimates (ASE) is the most popular criterion,^41, 57^ which assesses the difference between the predicted and actual model accuracies in a percentage scale:

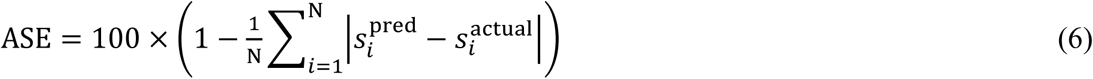

Where 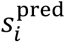 and 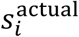 are the s-scores (eq. 5) of the predicted and actual Cα position errors on *i*^th^ residue, respectively. The ASE score ranges from 0 to 100, with higher score indicating more accurate predictor.

As shown in Figure 5, the ASE scores increase rapidly at the beginning of the simulations, and the ASE scores of predictions averaging five trajectories (MD average) are significantly better than that of each individual run. Compared with simulations at 330K, simulations at 380K give faster convergence of ASE scores during the trajectories. When the temperature is > 380K, the speed of convergence is similarly fast. At all temperatures, both ASE scores from MD average and individual MD runs can exceed ProQ3 in a very short time. At 380K, the ASE scores of MD average can exceed ProQ3D within 0.2 ns and reach near convergence within 0.4 ns. These results indicate that the performance of our MD-based method can be better than the state-of-the-art single-model MAE methods within a short simulation time.

**Figure 5.**
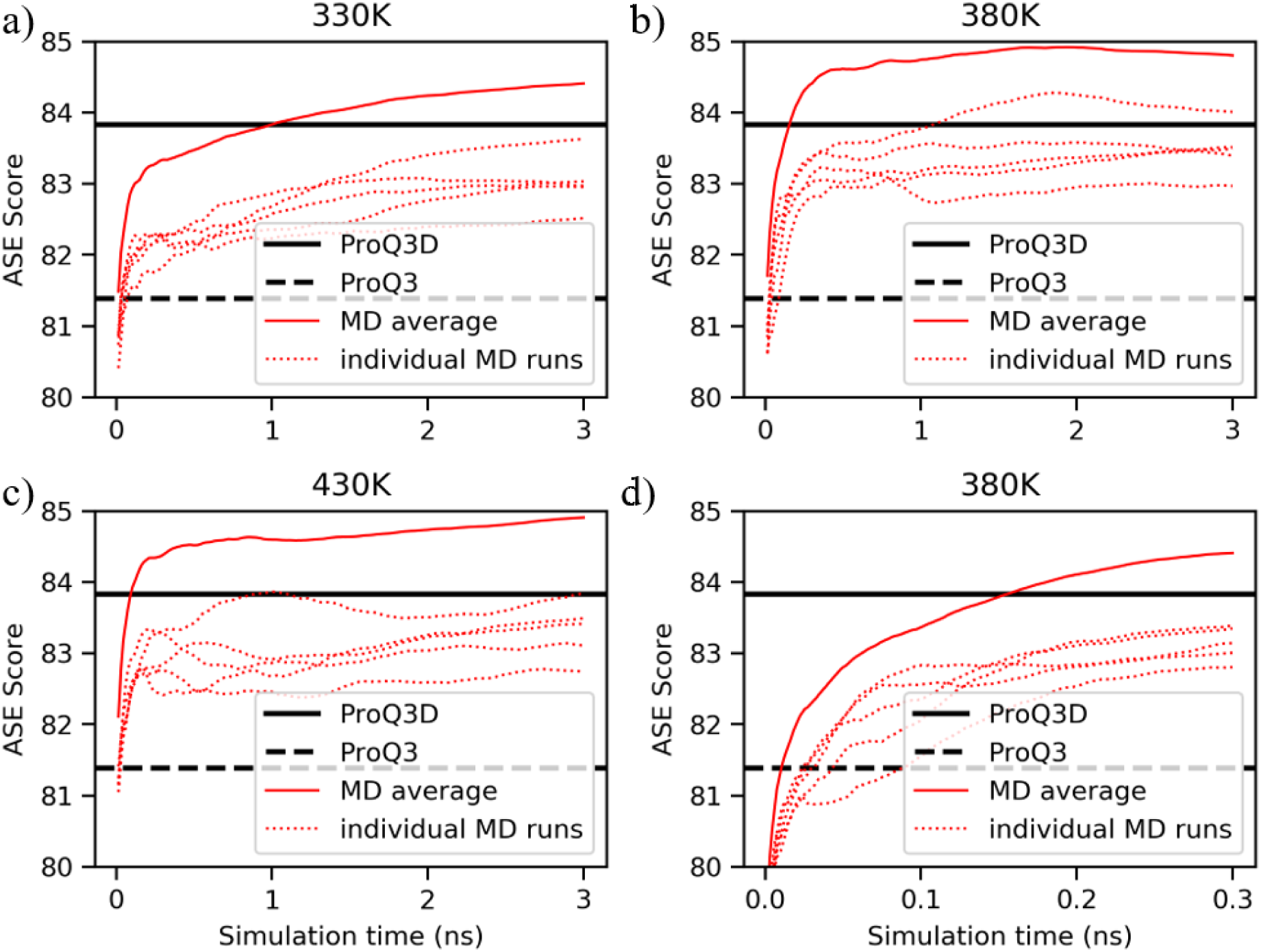
Average ASE scores (eq. 6) of MD predictions in the training dataset at a) 330K b) 380K c) 430K, over 3 ns simulation time. ASE scores of ProQ3 and ProQ3D were also shown for comparison. d) An enlarged view of the results from first 0.3 ns simulation in Figure 5b.

We also investigated whether our MD-based method can be used with force fields other than our RSFFs. Similar MD simulations were performed using a more popular AMBER force field ff99sb, which can be regarded as the parent of RSFF2. By averaging five parallel trajectories, ff99sb gave predictions better than ProQ3, and similar to ProQ3D when simulation time > 1 ns (Figure 6, Table S1&S2). However, the ff99sb predictions are still not as good as those from RSFF2, which is consistent with our observations in *ab initio* protein folding.^50^ For 20 out of 31 proteins, RSFF2 gives ASE scores higher than ff99sb. Thus, MD simulation can be used with other force fields for model accuracy estimation, but the RSFF2 force field may perform better.

**Figure 6.**
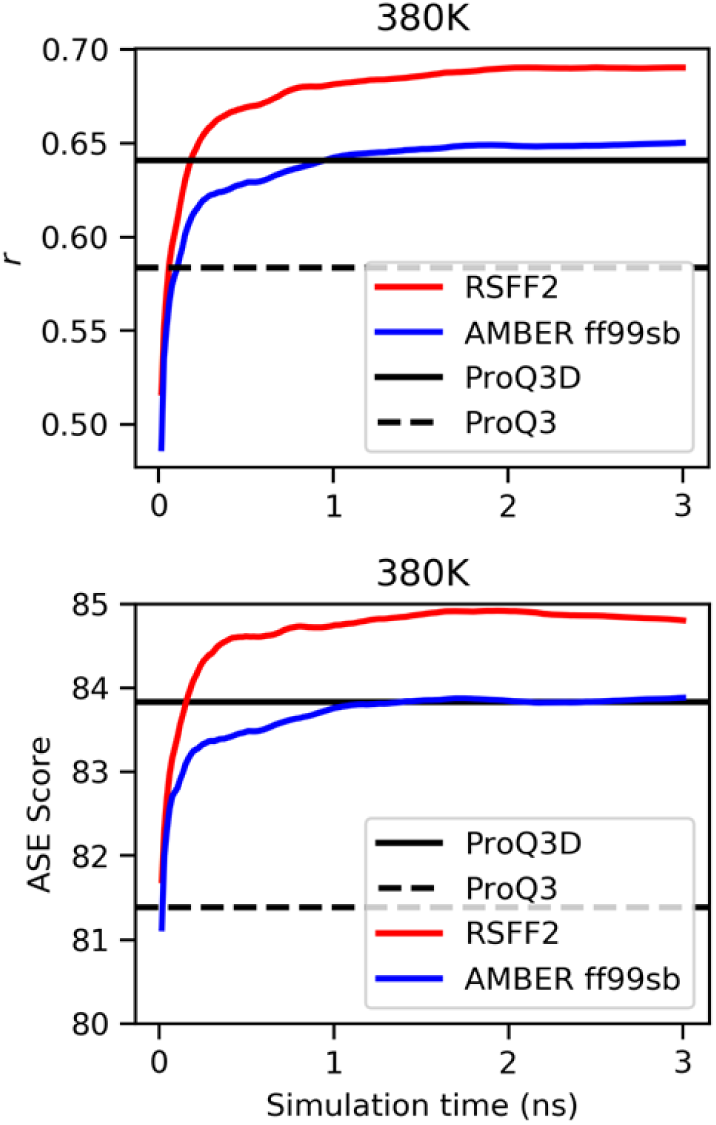
Upper: correlation coefficients (*r*) between MD predictions (averaging five parallel trajectories at 380K) and actual errors (in s-scores) over the 3 ns simulations, with a comparison between predictions from the RSFF2 force field and those from AMBER ff99sb. Lower: related ASE scores of the MD predictions. Results from ProQ3 and ProQ3D were also shown for comparison.

### Combining MD-based MAE predictor with ProQ3D

As shown in Figure 7a, ASE scores from MD-based predictions are > 80 for 27 out of 31 protein models and are > 90 for 6 protein models. On the other hand, ProQ3D gives ASE score > 90 for only one model. However, larger fluctuations of performances among different proteins can be observed for the MD-based method, with standard deviation of 6.34 compared with 5.08 from ProQ3D. We also found that, the correlation coefficient between MD-based predictions and ProQ3D predictions is 0.64, which is much smaller than the similarities among other MAE methods.^41^ Only in two proteins (TR568 and TR624), the scores of both methods are below 80. Considering that our MD-based approach is fundamentally different from previous MAE methods including ProQ3D, we speculated that there might be complementarity between them and, therefore, combined these two methods for more accurate estimation (as a meta-predictor):

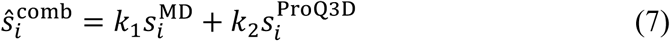

**Figure 7.**
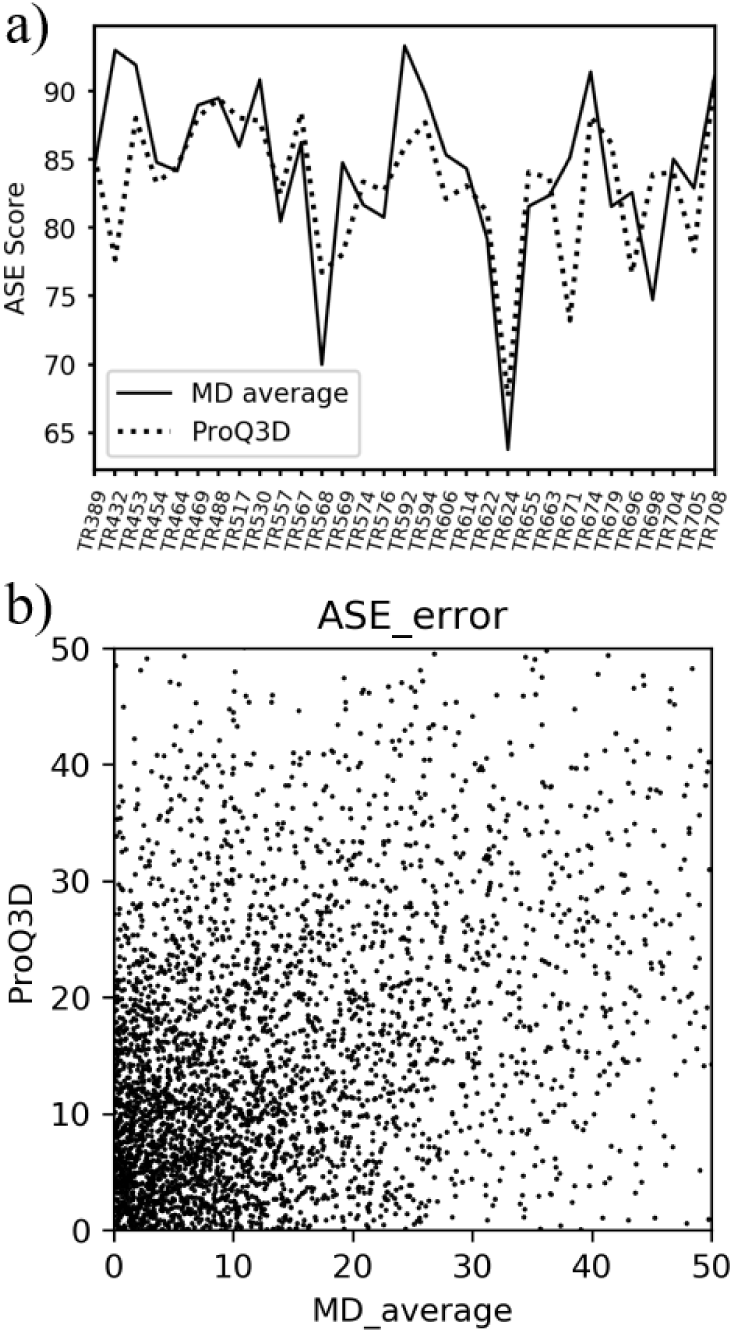
a) Average ASE scores of MD-based prediction (average of five parallel trajectories)

Figure 7. a) Average ASE scores of MD-based prediction (average of five parallel trajectories) and ProQ3D prediction for each protein model. b) Scatter plot of residue-wise ASE error (|predicted ASE score – actual ASE score|) of MD-based predictor and ProQ3D.

Consider the L1-norm penalty function in the ASE formula, least absolute deviations (LAD) regression (eq. 8) was used to optimize the coefficients (*k*_1_, *k*_2_) of MD-based and ProQ3D-based terms in eq. 7:

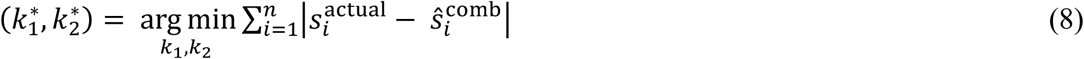

Figure 8 shows the optimized *k*_1_ and *k*_2_ as a function of simulation time *t*. When *t* < 0.2 ns, the contribution of MD-based predictor (*k*_1_) increases with simulation time, but keeps less than that of ProQ3D (*k*_2_). When *t* > 0.2 ns, the contribution of MD-based predictor becomes slightly larger than that of ProQ3D, and keeps near invariant with increasing *t*.

**Figure 8.**
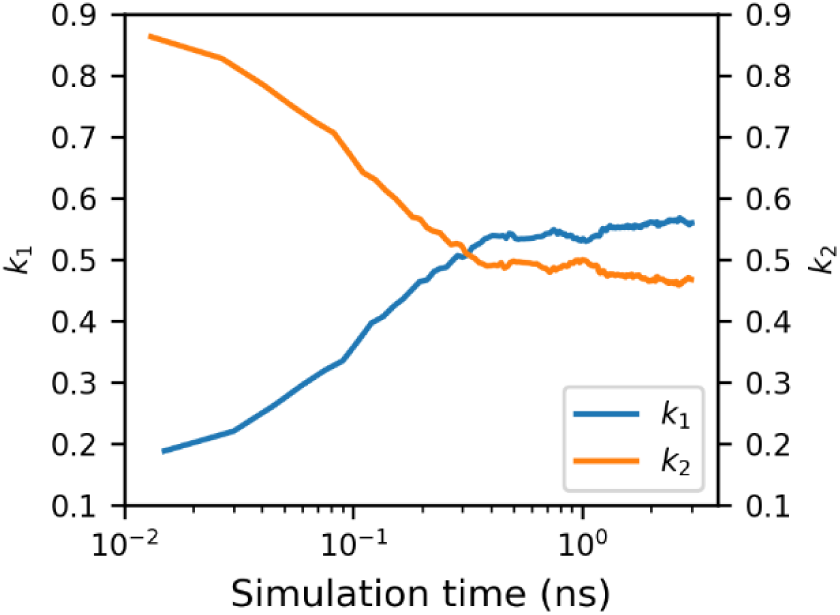
Combination coefficients *k*_1_ and *k*_2_ as a function of simulation time *t* (in logarithmic coordinates) at 380K.

As shown in Figure 9, the combined method (meta-predictor) performs significantly better than two parent methods. Combining with MD-based prediction from one individual trajectory of only 15 ps can improve ASE score of ProQ3D prediction by 0.7, which almost reaches the performance of MD average. When combining ProQ3D with MD average, the ASE score can reach up to 85.5, which is higher than the ASE scores of ProQ3D and MD average by 1.7 and 0.9, respectively. This performance show that a simple linear combination of our MD-based predictor and the ProQ3D predictor can achieve better performance than any of the constitute methods.

**Figure 9.**
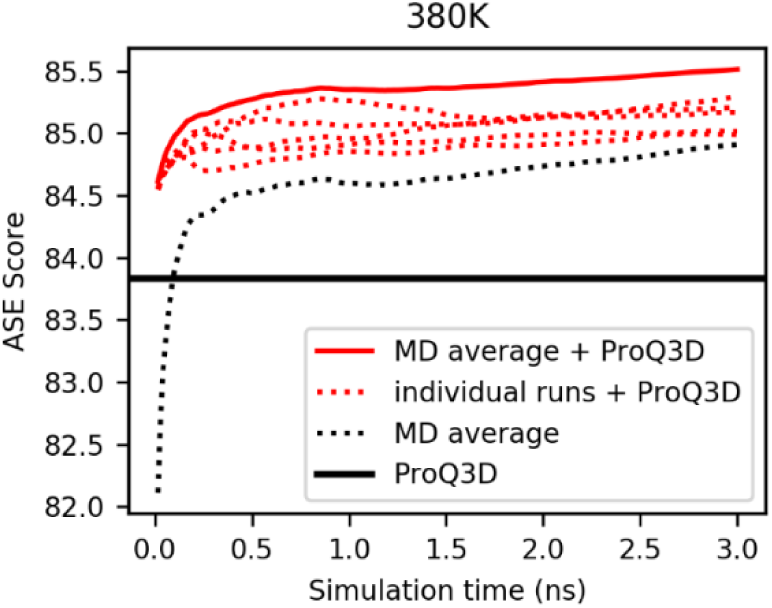
Average ASE scores of the combination (red lines) of MD-based method and ProQ3D in the training dataset compare with pure MD-based method (average, black dotted line) and ProQ3D (black solid line) over 3 ns simulation time at 380K.

**Figure 10.**
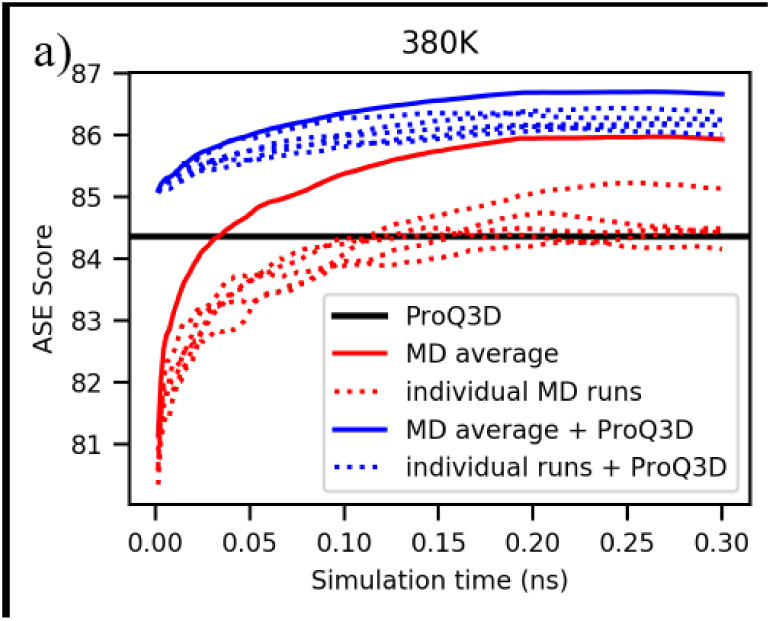
Average ASE scores of ProQ3D, MD-based predictor and the combination method from the test dataset. MD simulations were carried out at 380K for 0.3 ns.

### Validation of the MD-based methods on the test set

Although the MD-based method is physics-based without any optimized parameter, there are two parameters (*k* and *b* in eq. 3) based on the statistical analysis of a set of protein models (training set). Besides, for the meta-predictor combining MD and ProQ3D, there are two additional parameters (*k*_1_, *k*_2_) optimized on the training set. To validate the performance of our new methods on unseen data, 24 models of different proteins are chosen for testing. For each model, five parallel MD simulations were carried out at 380K for 0.3 ns using the RSFF2 force field. All the parameters are the same as above. As expected, MD predictions on the test set perform no worse than those on the training set. The ASE scores of MD-based predictor can exceed ProQ3D within 0.03 ns and reach 86 within 0.3 ns (Figure 9). In 17 out of 24 cases, MD-based predictor can give better performance than ProQ3D, and the ASE score of the meta-predictor combining MD average and ProQ3D can reach 86.7, which is higher than the ProQ3D prediction by 2.4. Our MD-based methods have good transferability on different proteins.

### Comparison with MD-based predictions from experimental structures

The positions of more flexible residues (such as those in loop) are in general more difficult to be predicted accurately in PSP, and the flexibility can be captured in MD simulation. To see whether our method can really predict residues’ position errors, we performed MD simulations of 31 target proteins starting from their experimental structures, under the same settings. The residue-wise RMSD values obtained in these simulations can be regarded as representing the intrinsic structural flexibility at residue level. The average residue-wise RMSD from simulations of experimental structures (1.16 Å) is significantly smaller than that from simulations of model structures (1.93 Å). The MD-predicted per-residue model qualities (s-scores) of experimental structures are almost distributed close to 1.0, much higher than those of model structures (Figure 11). Especially, MD-predicted s-scores of loop residues in model structures have considerable occurrences in a large range of 0.2∼1.0, whereas a vast majority of loop residues have s-cores > 0.7 in experimental structures. This indicates that our prediction method can really distinguish different modeling accuracies at residue level, rather than just rely on measuring the residue-wise fluctuations (flexibilities).

**Fig 11.**
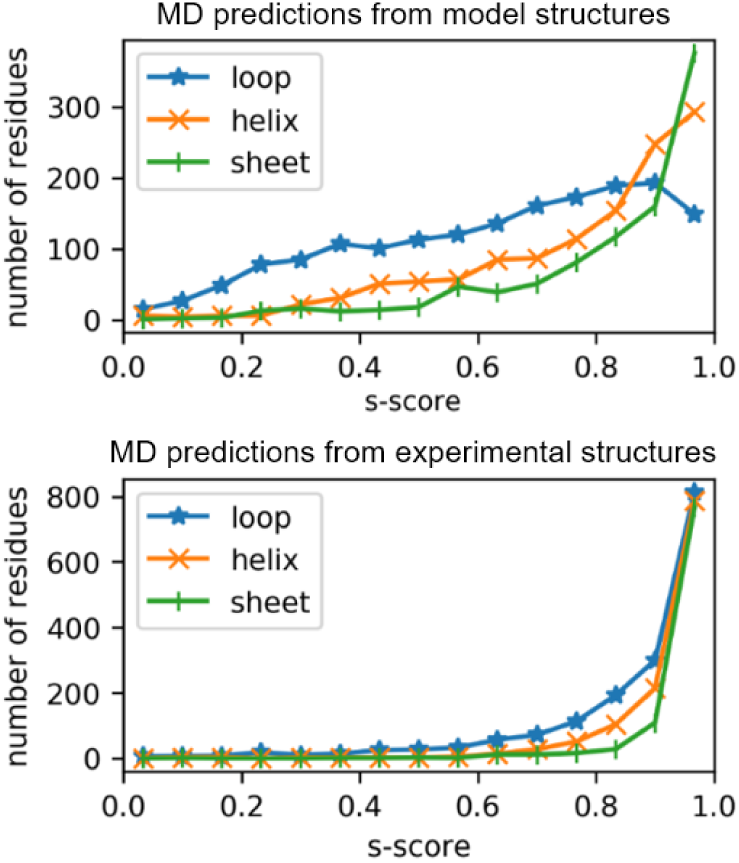
Distributions of predicted s-scores from MD simulations using (top) model structures and (bottom) experimental structures. Residues in different secondary structures (loop, helix, and sheet) were counted separately.

We found that the correlation coefficient *r* between the residue-wise flexibilities and the actual model errors is 0.44, much lower than the correlation achieved in our MD-based prediction (*r* = 0.68). We also performed linear regression between the intrinsic flexibilities and the MD-based predictions. The obtained residual can be regarded as predictions that does not contain any information about the intrinsic flexibilities. The correlation coefficient between the obtained residual and actual residue-wise error is 0.53. Therefore, if we excluded this information regarding intrinsic flexibility, MD-based method can still predict the modeling error of each residue, but in lower accuracy. Actually, prediction results for almost all other MAE methods have flexibility-related features such as predicted surface area. It has been proved that these features are important for better prediction of residue-level model accuracy.^26^

### Analysis of the MD trajectories

As shown in Figure 12a, we decomposed the movement of each residue in a MD trajectory into two directions, using a local (residue-specific) coordinate system after global superposition of the simulated structure and the corresponding experimental structure. The direction from each residue’s Cα position in the initial model to that in the experimental structure is defined as the “refinement axis” (X axis), because this should be the direction of an ideal model refinement. In addition to the refinement axis, residues can also move in a perpendicular direction (Y-axis). As shown in Figure 12b, most of the residues are located on the X > 0 side, indicating that they indeed move toward the corresponding positions in the experimental structures (positive refinement). Nevertheless, this tendency is not strong. In agreement with above findings, when the initial model is further away from the experimental structure, the movements on both the X and Y directions tend to be larger (Figure 12c). Noticeably, the movements towards negative X values (away from experiemental structure) also become larger for residues with larger errors, and when residues in the model are very far from the experimental positions, there is *not* a clear tendency for moving in the direction of better model quality. Thus, using short MD simulations, we can estimate the model accuracy even if good refinement is not achieved. In this work, we assumed that the relationship between residue-wise actual errors 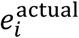 and time-averaged deviations in MD simulation 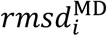 is linear (eq. 3). However, as we can see in Figure 12d, this relationship is actually weaker than linear. For residues with larger position errors, their movements in MD simulation related to the actual errors (|***v*_1_**|) are significantly smaller (in both X and Y directions).

**Figure 12.**
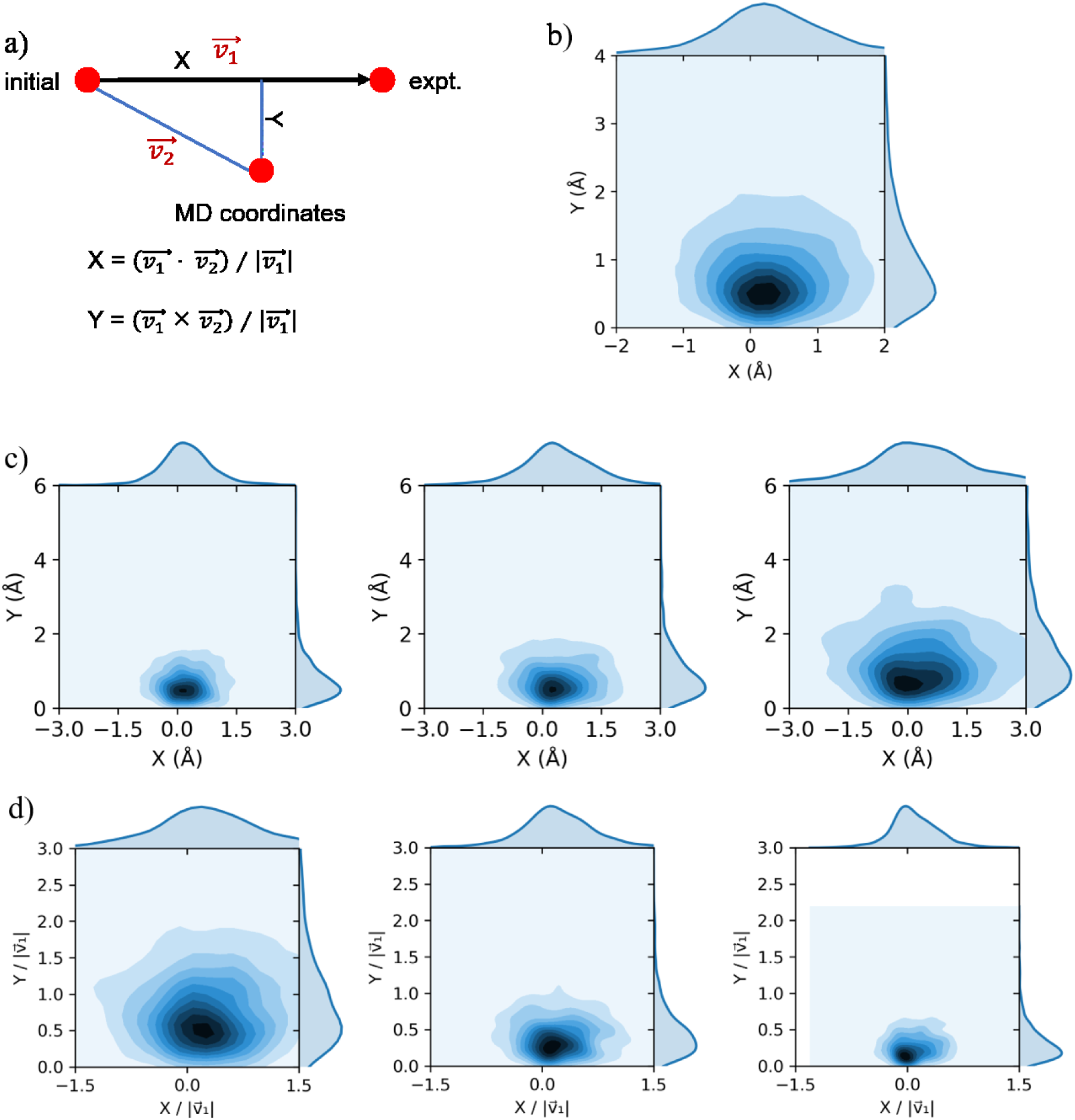
a) Local coordinates decomposition of residue-wise movements during MD simulations into X (refinement) axis and Y axis. b) X-Y distribution of all residues in the training dataset. c) X-Y distributions of residues in different ranges of actual residue-wise errors: small (0, 1.39 Å], medium (1.39 Å, 2.65 Å], and large (2.65 Å, 29.9 Å] from left to right. Each group has the same number of residues. d) X-Y distributions in (c) normalized by actual residue-wise errors. All MD simulations were carried out at 380K for 0.3 ns.

## Discussion

In principle, the relationship between the sequence of a protein and its native structure is governed by physical chemistry.^58^ Native structure of a protein corresponds to the global minimum on its free energy surface. Indeed, experimental structures of some small proteins can be predicted by simulating their spontaneous folding using molecular dynamics (MD).^59, 60^ However, the most successful protein structure prediction methods are usually based on data mining of known protein sequences and structures.^3, 13, 61, 62^ These knowledge-based methods are much more efficient than physics-based simulations, but they are less physically realistic, especially neglecting the solvent molecules which can play very important roles.^63-65^

Recently, with rapid developments of computational hardware and software, MD simulation has been attracting increasing applications in studying conformational behaviors of peptides and proteins. This has been motivating continuous improvements of protein force fields, along with increasing predicting power. Feig and coworkers showed that restrained MD simulations (a few hundred nanoseconds or less) in explicit water can result in improvements of structure models.^46, 47^ We have also shown that long-time replica-exchange MD simulations (microseconds per replica) sometimes can result in significant model improvements.^48^ In recent CASP competitions, most top-ranking groups in the refinement category use certain type of MD simulation.^49^ However, consistent improvement of structure models can only be achieved by applying restraints to keep structures close to the initial positions. This conservative strategy limits structural improvements that can be achieved. On the other hand, more extensive conformational sampling sometimes can give more significantly improved structures, but may also lead to much worse structures.^48^ If we can put more restraints on the good quality parts of a model while let less accurate parts more freely to move, more efficient conformational sampling and better refinement may be achieved.

In this work, we found that MD simulations can also be used to estimate model accuracy at residue level, with much less computational cost compared with structure refinement. During MD simulation, significant deviations from the initial position can be observed quickly (in tens of picoseconds), and residues with larger deviations from experimental structures indeed tend to move further away from their initial positions. However, from our analysis, there is not a strong overall trend for the positions of residues to move towards the corresponding native structures. Thus, it is difficult to use these trajectories to optimize the protein structures. This may be because these residues may take a more complicated transition path and time scale much longer than a few nanoseconds to achieve significant refinement. Therefore, they do not move in the right direction initially. However, for MAE, it seems that a series of short MD simulations can achieve good results. We also found that averaging parallel trajectories can achieve better predictions, this can be understood because a MD trajectory is a stochastic search in the conformational space. Further increasing the number of trajectories in our MD-based method may achieve better performance and more stable predictions.

Even the performance of current MD-based MAE method is not much better than top single-model MAE methods, developing a new type of methods is still important. Often, a meta-predictor combining different methods can outperform any of the constituent methods,^39, 42, 43, 45, 66^ Indeed, combining a MD-based MAE predictor with ProQ3D can achieve better performance. This can be understood considering the fundamental difference between our MD approach and those knowledge-based MAE methods.

The 31 model in the training set have overall Cα RMSD of 1∼8 Å to corresponding native structures. For each model, we calculated the correlation coefficient *r* between the predicted residue-wise error and the actual residue-wise error (higher *r* value indicates more accurate prediction). It is found that the obtained *r* values and the overall RMSDs of models has a correlation coefficient of −0.197. In the 31 models, 16 of them have overall RMSD < 3.5Å and their average *r* is 0.731, while the average *r* of the other 15 targets (RMSD > 3.5Å) is 0.663. In contrast, these two average *r* values from ProQ3D predictions are 0.678 and 0.712, respectively. Thus, MD-based method is more accurate for models with higher quality (small overall RMSD), while the ProQ3D method has the opposite trend, although this trend is very weak.

In the recent CASP competitions, best consensus-based methods and hybrid methods perform better than best single-model methods for both global and local MAE (MQA). Still, CASP has been continuously emphasizing the importance of single-model methods.^41^ First, a large number of predicted models which are required by consensus-based methods are not easily available in every-day practice. Moreover, single-model methods appear to be better than consensus-based methods in picking the best model from decoy sets, according to the results of recent CASP12.^41^ Finally, single-model methods can be combined with consensus-based methods to achieve a better performance.^66^ Therefore, the improvement of the single-model method still play a crucial role in the future development of better MAE methods.^27, 34^ Here we limit the scope of this work to “a proof of concept” study. However, in the future, more sophisticated methods will be developed, including those utilize the machine learning to combine various structural features from MD simulation, as well as traditional features for model quality assessment.

## ASSOCIATED CONTENT

## Supporting information

Correlation coefficients (Table S1) and ASE scores (Table S2) from ProQ3/ProQ3D predictions, as well as MD-based predictions using the RSFF2 and AMBER ff99sb force fields. The Supporting Information is available free of charge via the Internet at http://pubs.acs.org.

## Notes

The authors declare no competing financial interest.

## ACKNOWLEDGEMENTS

We thank the financial supports from the National Natural Science Foundation of China (21573009), and the Shenzhen Science and Technology Innovation Committee (JCYJ20170412150507046).

